# Perforin-2 Permeabilizes the Envelope of Phagocytosed Bacteria

**DOI:** 10.1101/274878

**Authors:** Fangfang Bai, Ryan M. McCormack, Suzanne Hower, Gregory V. Plano, Mathias G. Lichtenheld, George P. Munson

## Abstract

Perforin-2, the product of the *MPEG1* gene, limits the spread and dissemination of bacterial pathogens in vivo. It is highly expressed in murine and human phagocytes, and macrophages lacking Perforin-2 are compromised in their ability to kill phagocytosed bacteria. In this study we used *Salmonella typhimurium* as a model intracellular pathogen to elucidate the mechanism of Perforin-2‘s bactericidal activity. In vitro Perforin-2 was found to facilitate the degradation of antigens contained within the envelope of phagocytosed bacteria. In contrast, degradation of a representative surface antigen was found to be independent of Perforin-2. Consistent with our in vitro results a protease sensitive, periplasmic superoxide disumutase (SodCII) contributed to the virulence of *S. typhimurium* in Perforin-2 knockout but not wild-type mice. In aggregate our studies indicate that Perforin-2 breaches the envelope of phagocytosed bacteria facilitating the delivery of proteases and other antimicrobial effectors to sites within the bacterial envelope.

## Introduction

Macrophages and neutrophils phagocytose microorganisms to remove them from blood and tissues. As the phagosome matures the multisubunit NADPH oxidase assembles on its membrane and reduces O_2_ to generate superoxide in the lumen of the phagosome (Karimi et al., 2014; Nauseef, 2004). The products of this respiratory burst –superoxide and subsequently other reactive oxygen species (ROS)– are bactericidal. The destruction and degradation of phagocytosed microbes is further facilitated by acidification of the phagosome and fusion with lysosomes that deliver oxygen-independent antimicrobial effectors such as lysozyme, glycosylases, proteases, and other hydrolases (Cederlund et al., 2011). Large antimicrobials such as proteases and other hydrolases are typically membrane impermeable molecules. This property is advantageous in that it allows them to be confined within lysosomes and phagolysosomes. However it also precludes them from reaching the internal components of phagocytosed bacteria. For example, lysozyme hydrolyzes β(1,4)-glycosidic bonds of peptidoglycan; the primary structural component of bacterial cell walls. In gram-negative bacteria peptidoglycan resides in the space between the outer and inner membranes; i.e., the periplasm. Thus, for these bacteria a breach of the outer membrane must precede lysozyme-dependent degradation of peptidoglycan (Ellison and Giehl, 1991; Martinez and Carroll, 1980). For hydrolases with targets in the cytosol of gram-negative bacteria the challenge is two-fold as their substrates are bound by both an inner and outer membrane.

Studies over the past two decades have shown that antimicrobial peptides such as the defensins and cathelicidins attack and disrupt bacterial membranes (Gallo et al., 1997; Turner et al., 1998; Wiesner and Vilcinskas, 2010; Zanetti, 2004). NMR studies of the cathelicidin LL-37 suggest that the peptide destabilizes the bacterial membrane by carpeting rather than penetrating the lipid bilayer (Henzler Wildman et al., 2003). Within phagolysosomes the murine cathelicidin CRAMP has been shown to be active against phagocytosed *Salmonella*; most likely by disruption of the bacterium’s outer membrane (Kim et al., 2010; Rosenberger et al., 2004). Likewise, cathelicidin LL-37 may play a similar role in human phagocytes (Sonawane et al., 2011; Stephan et al., 2016). Nearly coincident with the initial descriptions of LL-37 and CRAMP the *Mpeg1* gene was identified as a potential marker of mammalian macrophages due to its relatively high expression in mature human and murine macrophages (Gallo et al., 1997; Gudmundsson et al., 1996; Spilsbury et al., 1995). *Mpeg1* encodes a 73 kDa protein referred to as Perforin-2 and in their initial report Spilsbury et al. noted its partial homology to the membrane attack complex perforin (MACPF) domain of Perforin; the cytolytic protein of natural killer cells and cytotoxic T lymphocytes (Spilsbury et al., 1995). Unlike the carpet mechanism of cathelicidins Perforin is a large polypeptide that polymerizes on target membranes. A concerted structural transition results in a pore through the lipid bilayer whose hydrophilic channel is lined with amphipathic β-strands donated by the MACPF domains (Law et al., 2010; Voskoboinik et al., 2015). It is through this channel, either at the cell surface as originally hypothesized or within endosomal membranes as a more recent study suggest, that granzyme proteases enter tumor and virally infected cells to facilitate their destruction and lysis (Lichtenheld et al., 1988; Podack et al., 1989; Thiery et al., 2011). MACPF domains are also present in the terminal complement proteins which form pores in the outer membranes of gram-negative bacteria through a similar mechanism of polymerization and structural transition (Dudkina et al., 2016).

Despite the homology of mammalian Perforin-2 to known pore forming proteins there was no further elaboration of its function until 2013; nearly two decades after the initial report of Spilsbury et al.(McCormack et al., 2013; Spilsbury et al., 1995). In 2013 McCormack et al. demonstrated that the expression of Perforin-2 correlated with the killing of phagocytosed gram-negative, - positive, and acid-fast bacteria in vitro (McCormack et al., 2013). Subsequent studies with transgenic mice found that Perforin-2 -/- mice are unable to limit the proliferation and dissemination of infectious bacteria. Not surprisingly, Perforin-2 knockout mice succumb to infectious doses that are non-lethal to their wild-type littermates (McCormack et al., 2016; McCormack et al., 2015a; McCormack et al., 2015b). Moreover this defect is not limited to a particular route of infection nor pathogen. Nor is it limited to mammalian Perforin-2 as similar results have been reported with zebrafish as the model organism (Benard et al., 2015). In aggregate these latter studies have demonstrated that Perforin-2 is associated with broad spectrum bactericidal activity. In this study we utilized *Salmonella typhimurium* as a model pathogen to elucidate the mechanism of Perforin-2 dependent killing of phagocytosed bacteria.

## Results

### Perforin-2 limits the survival of phagocytosed bacteria independent of ROS

Because the interactions between *Salmonella* sp. and macrophages have been extensively characterized, we chose to exploit the *Salmonella*/phagocyte paradigm to probe the mechanism of Perforin-2 dependent killing of phagocytosed bacteria (Steele-Mortimer, 2008). Accordingly, peritoneal exudate macrophages (PEMs) and neutrophils isolated from wild-type and Perforin-2 - /- mice were infected with *Salmonella enterica* serovar Typhimurium (hereafter *S. typhimurium)*. As expected from previous studies that have shown *Salmonella* survives within macrophages, the intracellular load of bacteria either increased or remained constant in wild-type phagocytes (Figure 1AB). However, Perforin-2 deficient phagocytes had significantly higher intracellular loads of *S. typhimurium* than wild-type phagocytes (Figure 1AB). This demonstrates that Perforin-2 limits the survival and/or replication of phagocytosed *S. typhimurium* and is consistent with previous studies that have shown Perforin-2 is a potent antimicrobial effector against *Salmonella* as well as gram-positive and acid-fast bacteria (Fields et al., 2013; McCormack et al., 2016; McCormack et al., 2015a; McCormack et al., 2015b).

**Figure 1.**
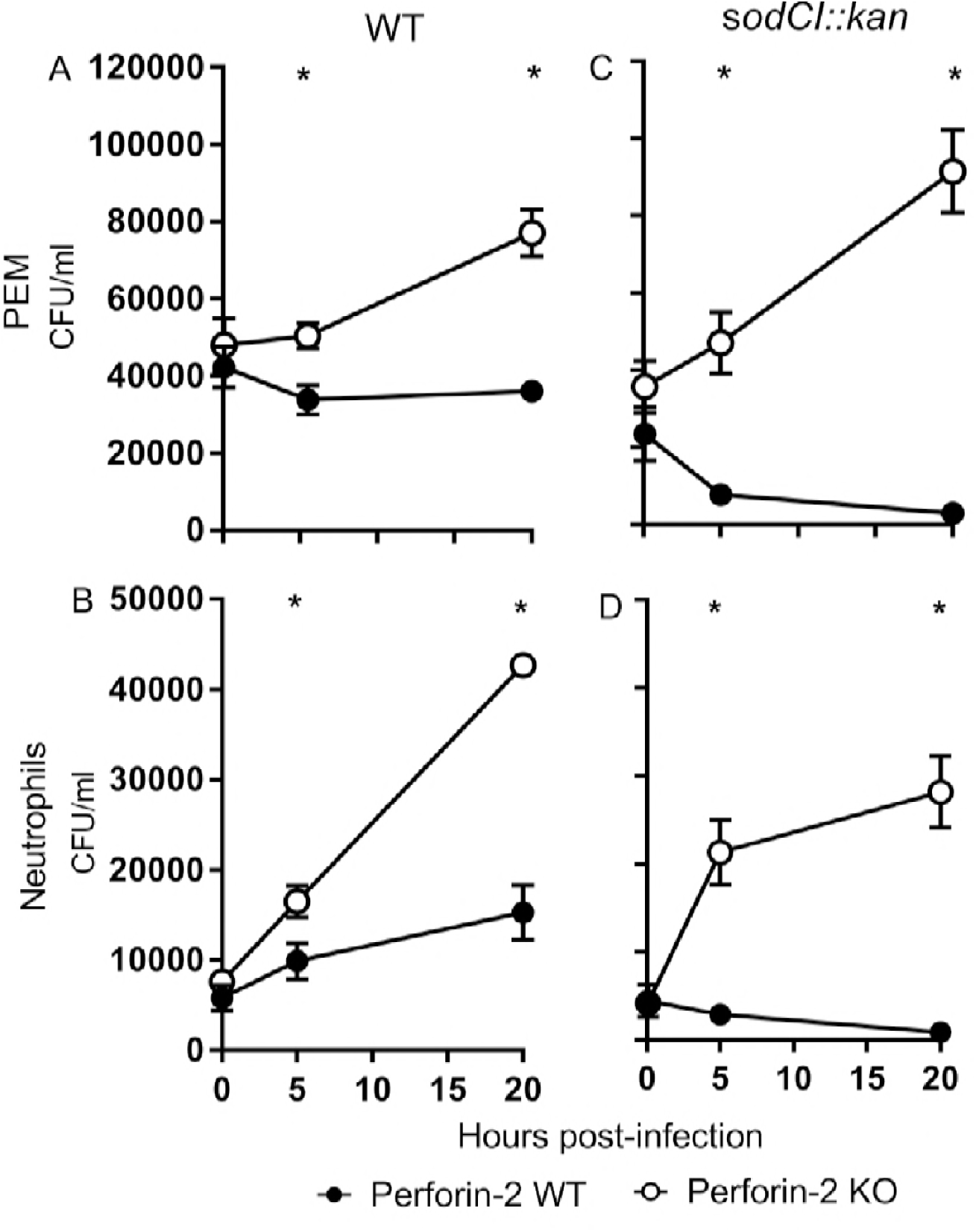
Perforin-2 limits the survival of phagocytosed bacteria independent of ROS. As indicated PEM or peritoneal neutrophils were isolated from wild-type and Perforin-2 -/- (KO) mice and stimulated with IFN-γ for 14 hours prior to infection with (AB) wild-type *S. typhimurium* strain GPM2004, or (CD) strain ST188 [*sodCI::kan*]. Gentamicin was used to eliminate extracellular bacteria and intracellular bacteria were enumerated by plating cellular lysates. Means and standard deviations are shown; *n* = 3. \**P* ≤ 0.05, Student’s *t* test

One of the aforementioned studies also investigated the relationship between reactive oxygen species (ROS) and Perforin-2 and concluded that the bactericidal activity of ROS was dependent upon Perforin-2 (McCormack et al., 2015a). This was based on two principle findings. First, chemical inhibition of ROS production significantly enhanced the survival of phagocytosed *S. typhimurium* –relative to mock treated cells– in wild-type but not Perforin-2 deficient PEMs. Second, wild-type PEMs killed a Δ*sodCI* strain of *S. typhimurium* much more efficiently than wild-type *S. typhimurium*. SodCI is a periplasmic superoxide dismutase that neutralizes ROS and thus promotes the survival of *S. typhimurium* within phagosomes (Krishnakumar et al., 2004; McCormack et al., 2015a). However, the *sodCI* mutant was found to be no less fit than the wild-type strain when Perforin-2 -/- PEMs were used. Thus, both a chemical and genetic analysis suggest that ROS is not a significant bactericidal effector when Perforin-2 is absent.

As with the previous study we observed similar effects with both PEMs and neutrophils. In either case the *S. typhimurium sodCI* mutant was efficiently killed by Peforin-2 proficient but not deficient phagocytes (Figure 1CD). The source of phagocytic ROS is the multisubunit NADPH oxidase NOX2 (Panday et al., 2015). Assembly of the enzymatic complex involves the translocation of cytosolic proteins to the endosomal membrane to form the active complex that generates the respiratory burst (Panday et al., 2015). Likewise Perforin-2, a transmembrane protein of cytosolic vesicles, dynamically translocates to and fuses with phagocytic vesicles containing bacteria (McCormack et al., 2015a; McCormack et al., 2015b). This raised the possibility that Perforin-2 is involved in the assembly and/or activation of the NADPH oxidase. If true the respiratory burst would be deficient in Perforin-2 -/- phagocytes and would account for the survival of *S. typhimurium sodCI* mutants in Perforin-2 deficient phagocytes.

To determine whether or not Perforin-2 is required for ROS production a luminol based chemiluminescence assay was used to quantify ROS productions in IFN-γ primed PEMs, peritoneal macrophages isolated without thioglycollate stimulation, and neutrophils from wild-type and Perforin-2 knockout mice. We found that ROS production was equally robust in wild-type and Perforin-2 -/- macrophages that were stimulated with PMA or LPS (Figure 2AB). The kinetics of ROS production was also similar as macrophages of both genotypes exhibited peak ROS production at 60 min. To confirm that the chemiluminescent signal was due to ROS production by a NADPH oxidase, phagocytes were pretreated with diphenyleneiodonium chloride (DPI); an inhibitor of NAPDH oxidases. As expected DPI treated cells produced negligible amounts of ROS (Figure 2). Likewise, ROS production was also negligible in unstimulated macrophages (data not shown). As with macrophages, we found few statistically significant differences between ROS production in wild-type and Perforin-2 -/- neutrophils (Figure 2CD). Thus, we conclude that impaired ROS production cannot account for the survival of *S. typhimurium* Δ*sodCI* mutants in Perforin-2 -/- phagocytes.

**Figure 2.**
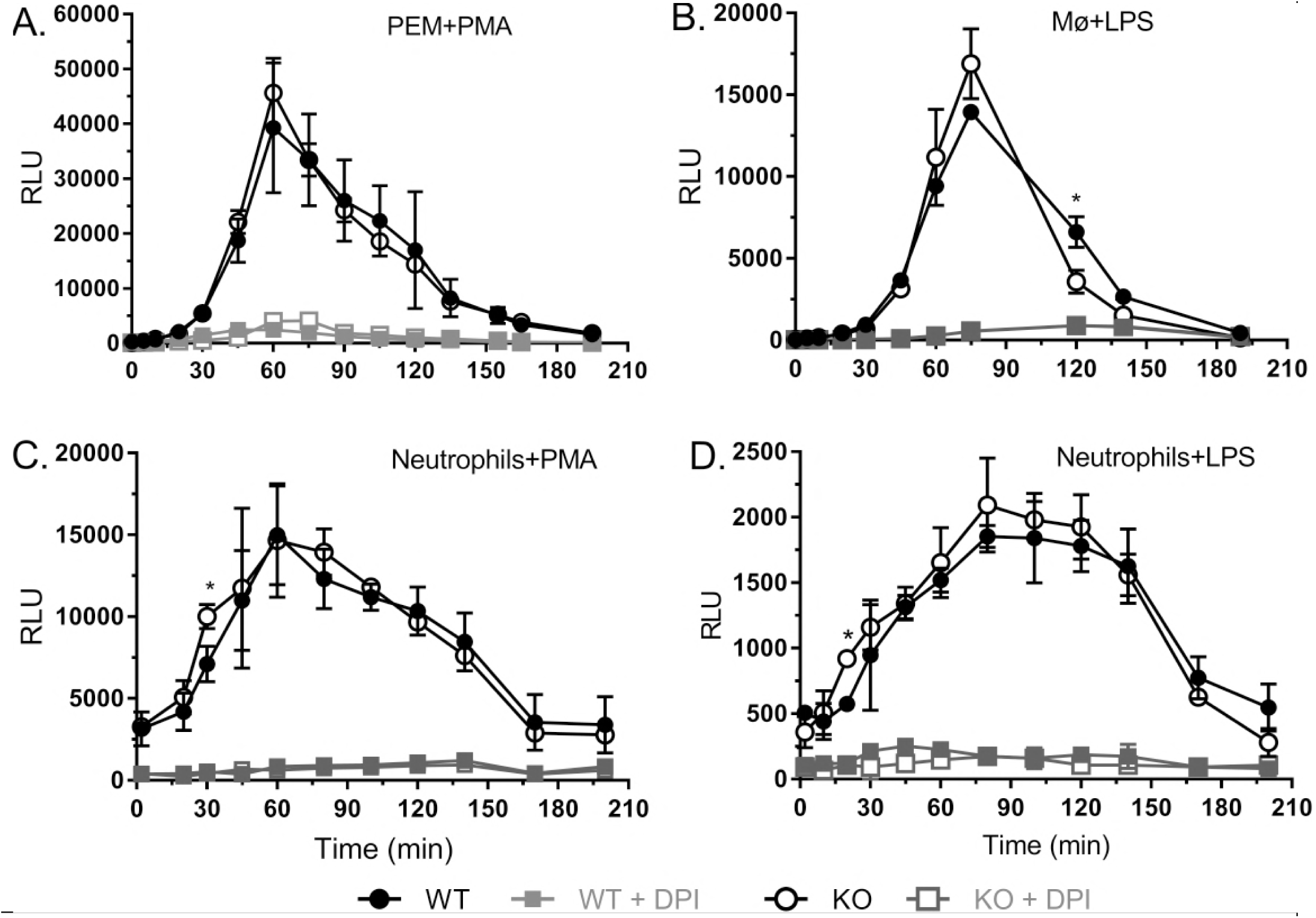
Perforin-2 is not required for ROS production in phagocytes. As indicated wild-type and Perforin-2 -/- (KO) peritoneal (A,B) macrophages and (C,D) neutrophils were stimulated with PMA or LPS to elicit ROS production which was detected by luminal based chemiluminescence. An enhancer was added to amplify chemiluminescence when macrophages were used. As indicated some cells were also treated with DPI, an inhibitor of the phagocytic NAPDH oxidase. ROS activity is reported as mean relative light units (RLUs) ± SD; *n* = 3. \**P* ≤ 0.05, Student’s *t* test.

### SodCI and SodCII are functionally redundant in Perforin-2 -/- phagocytes

Having excluded the possibility that Perforin-2 deficiency results in impaired ROS production, we next considered the possibility that *S. typhimurium sodCI* mutants are resistant to ROS in Perforin-2 deficient but not proficient phagocytes. As the genome of *S. typhimurium* encodes a second periplasmic superoxide dismutase (SodCII) we considered the possibility that it provides resistance to ROS in Perforin-2 -/- phagocytes even though previous studies have concluded that SodCII does not attenuate ROS toxicity in wild-type cells and animals (Kim et al., 2010; Krishnakumar et al., 2004). To determine whether or not SodCI and SodCII are functionally redundant we infected Perforin-2 proficient and deficient phagocytes with a *sodCI sodCII* double mutant. Unlike the Δ*sodCI sodCII*^+^ strain which was killed by wild-type but not Perforin-2 knockout phagocytes (Figure 1CD), the Δ*sodCI sodCII::kan* strain was killed by both Perforin-2 +/+ and -/- phagocytes (Figure 3AB). Moreover, treatment with the NADPH oxidase inhibitor DPI demonstrated that killing of the double mutant was ROS dependent in both Perforin-2 +/+ and -/- PEMs (Figure 4). Complementation of the double mutant with *sodCII* allowed the complemented strain to proliferate in Perforin-2 -/- but not wild-type phagocytes (Figure 3CD). Thus, in agreement with previous studies we conclude that SodCII does not protect against ROS in wild-type phagocytes. However, the nullification of SodCII is clearly dependent upon Perforin-2 because SodCII is able to protect phagocytosed *S. typhimurium* from the bactericidal effects of ROS in Perforin-2 -/- phagocytes.

**Figure 3.**
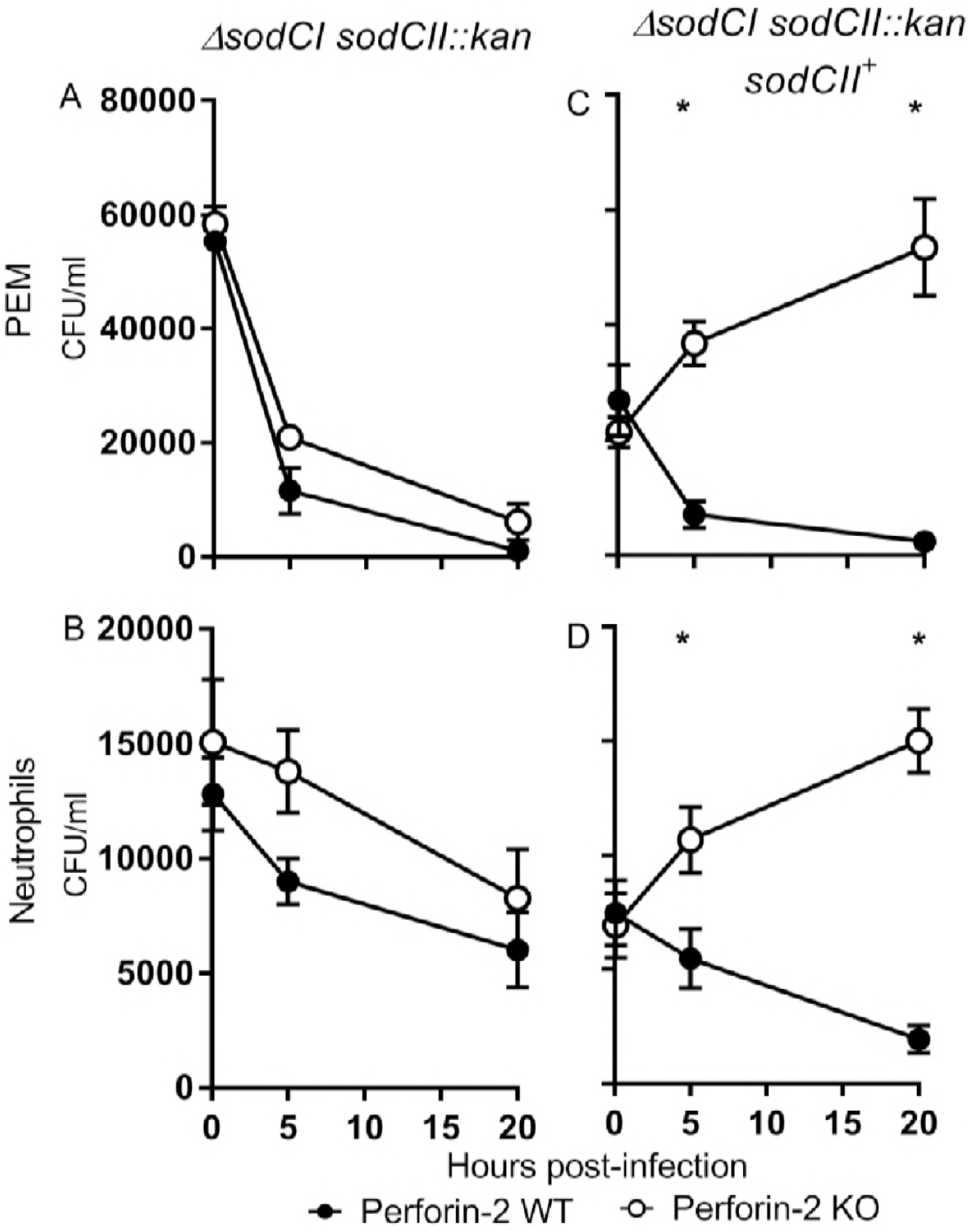
SodCII is functional in the absence of Perforin-2. As indicated PEM or peritoneal neutrophils were isolated from wild-type and Perforin-2 -/- (KO) mice and stimulated with IFN-γ for 14 hours prior to infection with *S. typhimurium* strain (AB) ST189 [Δ*sodCI sodCII::kan*] or (CD) GPM2008 [Δ*sodCI* Δ*sodCII::kan sodCII*^*+*^]. Gentamicin was used to eliminate extracellular bacteria and intracellular bacteria were enumerated by plating cellular lysates. Means and standard deviations are shown; *n* = 3. \**P* ≤ 0.05, Student’s *t* test.

**Figure 4.**
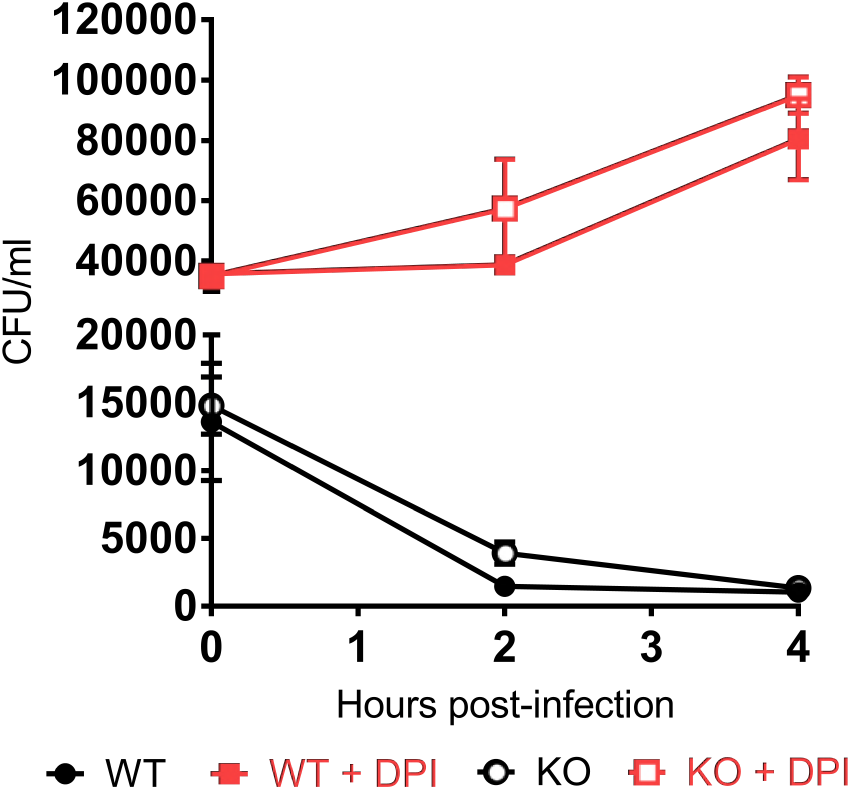
Inhibition of ROS production allows a Δ*sodCI* Δ*sodCII* mutant to proliferate intracellularly. PEMs from wild-type and Perforin-2 -/- (KO) mice were stimulated with IFN-γ 14 h prior to infection. As indicated some cells were also treated with DPI 30 min before infection with *S.* typhimurium strain ST189 [Δ*sodCI sodCII::kan*]. Gentamicin was used to eliminate extracellular bacteria and intracellular bacteria were enumerated by plating cellular lysates. Means and standard deviations are shown; *n* = 3.

### Perforin-2 facilitates the degradation of antigens enclosed within the bacterial envelope

Although SodCI and SodCII have similar enzymatic properties, only SodCI provides resistance to endosomal ROS in wild-type phagocytes (Krishnakumar et al., 2004). Studies by the Slauch laboratory have shown that this phenomenon is not the result of differential expression (Krishnakumar et al., 2004). Rather it is due to differential degradation of the two superoxide dismutases; SodCI is protease resistant while SodCII is protease sensitive (Kim et al., 2010; Krishnakumar et al., 2004; Krishnakumar et al., 2007). Thus, SodCII does not protect phagocytosed *S. typhimurium* because it is proteolytically degraded. However SodCII is functional in Perforin-2 -/- phagocytes (Figure 3). Therefore, we considered it possible that the degradation of SodCII is Perforin-2 dependent. To determine whether or not this is the case we infected wild-type and knockout PEMs with a strain of *S. typhimurium* expressing SodCII-FLAG so that we could track the persistence of SodCII in the periplasm with a monoclonal antibody against the FLAG epitope. We also used antibodies against DnaK and flagellin to monitor the abundance of cytoplasmic and surface antigens respectively. Western blots of intracellular bacteria recovered 18 hours after infection revealed that extracellular flagellin, which was abundant on the surface of the bacteria prior to phagocytosis, was efficiently degraded in both wild-type and knockout PEMs (Figure 5A). In contrast there was a clear difference in the abundance of SodCII in bacteria recovered from Perforin-2 -/- and +/+ PEMs (Figure 5A). This was not due to differences in bacterial load because the difference in SodCII abundance was statistically significant when experimental replicates were quantified and normalized to DnaK of the bacterial cytoplasm (Figure 5B). A further indication that the bacterial loads were similar in these experiments is the fact that the difference in the amount of DnaK normalized to host β-actin was insignificant while the difference in the amount of SodCII normalized to β-actin was significant (Figure 5B).

**Figure 5.**
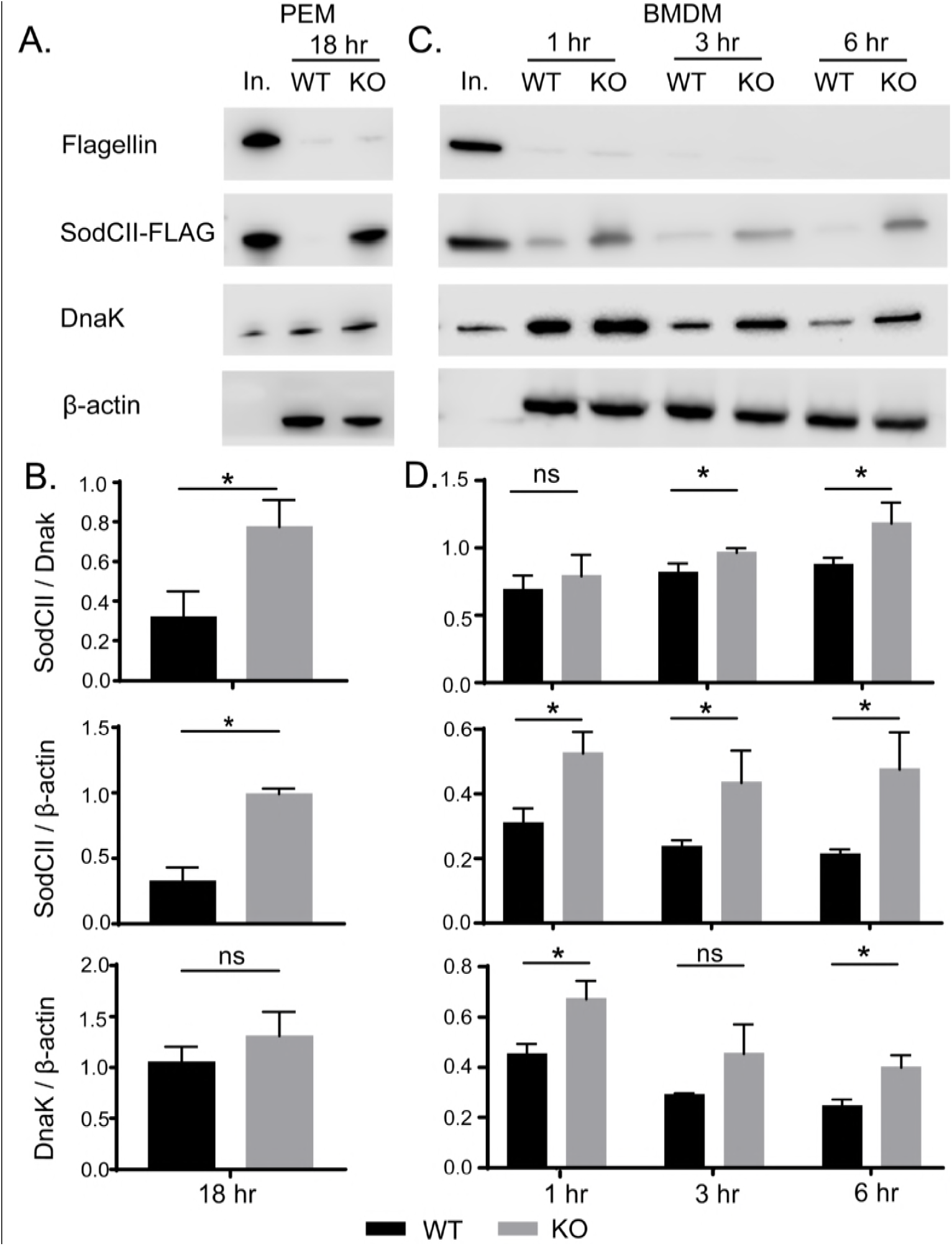
Perforin-2 facilitates the degradation of antigens enclosed within the bacterial envelope. (A)PEMs isolated from Perforin-2 +/+ and -/- (KO) mice were infected with a strain of *S. typhimurium* expressing SodCII-FLAG. After 18 hours the phagocytosed bacteria were recovered and the indicated antigens were detected by Western blot. (B)Quantification of Western blot data from three separate experiments in PEMs (C)Similar infection experiments were conducted with BMDM except the phagocytosed bacteria were recovered at 1, 3, and 6 hours post phagocytosis. (D)Quantification of Western blot data from three separate experiments in BMDM. Means and standard deviations are shown; *n* = 3. \**P* ≤ 0.05, Student’s *t* test. Abbreviations: In., inoculum; ns, not significant

We also examined the degradation of the three bacterial antigens at earlier time points. Because these experiments required significantly more phagocytes we used BMDM which can be obtained at higher yields than PEMs. As with PEMs extracellular flagellin was efficiently degraded in both wild-type and Perforin-2 -/- BMDM (Figure 5C). There was also apparent degradation of both SodCII and DnaK in both types of cells. Nevertheless there was less SodCII in bacteria recovered from wild-type than knockout phagocytes. This difference was statistically significant at 3 and 6 hours when normalized to DnaK even though concurrent degradation of DnaK likely results in an underestimation of the difference (Figure 5D). The degradation of DnaK also appears to lag that of SodCII; although, it is unclear if this is due to differences in protease accessibility and/or susceptibility. Differences in expression may also contribute to the apparent differences in degradation rates of SodCII compared to DnaK. Relative to β-actin the differences in SodCII in wild-type compared to Perforin-2 -/- BMDM was significant even at the 1 hour time point. This cannot be due to higher loads of the bacteria in Perforin-2 -/- phagocytes because the amount of DnaK normalized to β-actin also decreased over time; even in knockout cells. Additionally, extrapolation of our PEM bactericidal assays suggest the differences in bacterial loads is likely to be negligible at early time points; especially, at 1 hour. In aggregate these results demonstrate that Perforin-2 facilitates the degradation of internal –but not extracellular– antigens of phagocytosed bacteria.

### Perforin-2 negates SodCII in vivo

Having established that Perforin-2 facilitates the degradation of SodCII in vitro, the relevance of our observations were evaluated in a murine infection model. In brief, wild-type and Perforin-2 -/- mice were inoculated intraperitoneally with a mixture of wild-type *S. typhimurium* and a *sodCI* mutant at a 1:1 ratio. Four days after infection liver and spleen homogenates were plated on selective media to enumerate the load of each strain. Consistent with previous studies fewer *sodCI::kan* bacteria were recovered than wild-type bacteria recovered from Perforin-2 +/+ mice. The derived competitive indices were accordingly low and demonstrate that the *sodCI* mutant is significantly attenuated in Perforin-2 proficient mice (Figure 6A). In contrast, there was little to no attenuation of the *sodCI::kan* strain in Perforin-2 -/- mice as indicated by competitive indices near or at 1.0 (Figure 6A). Similar results were obtained with a strain of S. typhimurium that had spontaneously lost the pSLT virulence plasmid (Figure S1) (McClelland et al., 2001). Thus, SodCI is not essential in the absence of Perforin-2.

**Figure 6.**
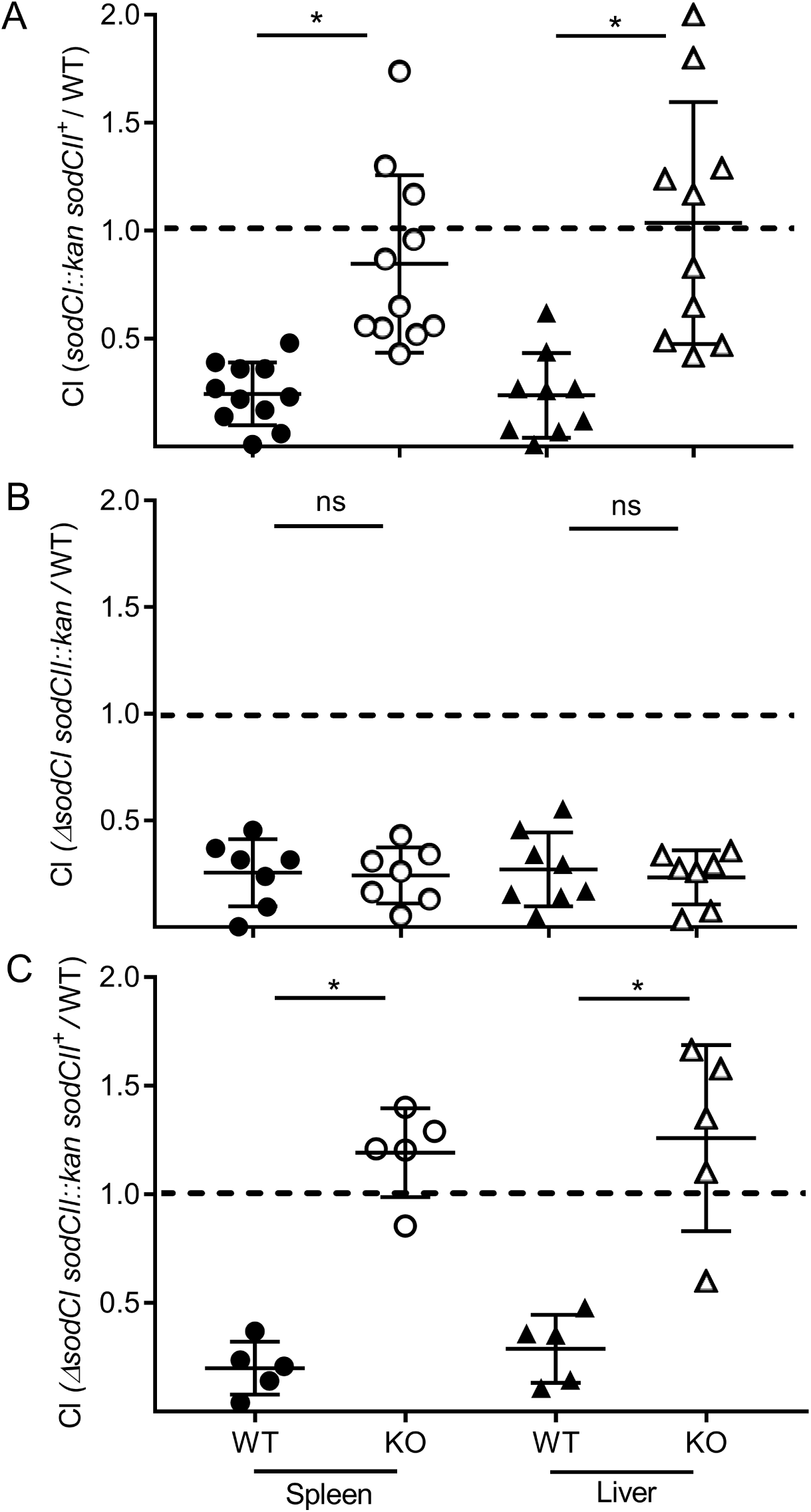
SodCII is functional in Perforin-2 knockout but not wild-type mice. Wild-type (WT) and Perforin-2 -/- (KO) mice were inoculated by i.p. injection with *S. typhimurium* wild-type strain GPM2004 and (A) Δ*sodCI::kan* strain ST188, (B) Δ*sodCI* Δ*sodCII::kan* strain ST189 or (C) Δ*sodCI* Δ*sodCII::kan sodCII*^*+*^ strain GPM2008 at a 1:1 ratio. Organs were harvested 4 days post infection and strains enumerated on selective media. Competitive indices are derived from the ratios of mutant strains to wild-type strains with compensation for any differences in inocula. Medians and standard deviations are indicated by horizontal bars. \**P* ≤ 0.05, *n* = 5-11. Student’s *t* test.

Based on our in vitro studies the persistence of SodCII was the most likely explanation for the lack of attenuation of the *sodCI* mutant in Perforin-2 knockout mice. Indeed this was found to be the case because a *sodCI sodCII* double mutant was found to be significantly attenuated relative to wild-type bacteria in Perforin-2 -/- mice (Figure 6B). Furthermore, complementation of the *sodCI sodCII* double mutant with *sodCII* resulted in a strain that was as virulent as wild-type *S. typhimurium* in Perforin-2 -/- mice (Figure 6C). In fact there appeared to be a slight competitive advantage of the complemented strain over the wild-type strain as indicated by competitive indices > 1. This could be the result of higher expression levels of *sodCII* from a heterologous promoter in our construct. In contrast, complementation with *sodCII* failed to rescue the double mutant in wild-type mice (Figure 6C). In aggregate the in vivo studies demonstrate that SodCII confers a protective advantage in Perforin-2 -/- but not wild-type mice. As such they are consistent with our in vitro finding that Perforin-2 facilitates the degradation of SodCII.

## Discussion

Perforin-2 is a type I transmembrane protein that we have previously shown localizes to endosomal vesicles as well as the endoplasmic reticulum, Golgi, and plasma membrane (McCormack et al., 2015a). A separate study that focused on the proteome of endocytic vesicles also found that Perforin-2 is present in endosomes following phagocytosis of latex beads by J774 macrophages (Duclos et al., 2011). The same study also found that Perforin-2 was more abundant in late endosomes and lysosomes than early endosomes. Perforin-2 is also coincident with subunits of the phagocytic NAPDH oxidase, proton transporters, and many other antimicrobial effectors of phagosomes and/or the phagolysosomes (Duclos et al., 2011; Nakamura et al., 2014). LPS stimulation of BMDM has also been shown to increase the abundance of Perforin-2 in endolysosomes compared to untreated cells (Nakamura et al., 2014). This is consistent with our own studies in which we reported that LPS results in the accumulation of Perforin-2 in vesicular structures and that Perforin-2 colocalizes with phagocytosed bacteria such as *Escherichia coli* and *S. typhimurium* (McCormack et al., 2015a; McCormack et al., 2015b). In aggregate these studies demonstrate that the subcellular distribution of Perforin-2 is consistent with its ability to facilitate the destruction of phagocytosed bacteria.

Although most phagocytosed bacteria are rapidly killed some are able to resist phagocytic antimicrobials and even survive within professional phagocytes. The latter includes *S*. *typhimurium* which must survive the respiratory burst and other antimicrobial assaults prior to the formation of *Salmonella* containing vacuoles; specialized niches within macrophages that afford the pathogen a more favorable environment than phagosomes or phagolysosomes (Anderson and Kendall, 2017). A central player in the survival of the respiratory burst is SodCI; a periplasmic superoxide dismutase that converts superoxide to hydrogen peroxide which is subsequently detoxified by bacterial catalases and peroxidases (Aussel et al., 2011; Hebrard et al., 2009; Slauch, 2011; Storz and Imlay, 1999). The pivotal role of SodCI in protecting *Salmonella* from ROS has been confirmed by several studies that have shown that *sodCI* mutants are more susceptible to ROS killing in vitro and less virulent than wild-type *Salmonella* in vivo (Craig and Slauch, 2009; Fang et al., 1999; Krishnakumar et al., 2004; Krishnakumar et al., 2007). However we have found that a *sodCI* null mutant is able to proliferate in Perforin-2 deficient phagocytes. Moreover the *sodCI* null mutant is as virulent as wild-type *S. typhimurium* in Perforin-2 deficient – but not proficient– mice. Because we have found that the production of phagocytic ROS is independent of Perforin-2, the survival of the *sodCI* mutant is not due to differences in ROS production. Rather it is due to the persistence of SodCII, a second periplasmic superoxide disumutase, in Perforin-2 deficient phagocytes. Consistent with our in vitro results we have also found that SodCI and SodCII are functionally redundant in Perforin-2 knockout mice.

The persistence of SodCII was unexpected because previous studies have shown that it is normally degraded by proteases of the phagolysosome (Krishnakumar et al., 2004; Krishnakumar et al., 2007). In contrast SodCI is resistant to proteolytic degradation and thus provides resistance to ROS in the phagolysosome (Krishnakumar et al., 2004). Our results in Perforin-2 +/+ cells and animals are consistent with these latter studies because *sodCII* is unable to complement a *sodCI sodCII* double mutant. However, *sodCII* is able to complement the double mutant in Perforin-2 -/- animals and isolated phagocytes. Thus, the presence of Perforin-2 is associated with the inactivation of SodCII.

How does Perforin-2 inactivate SodCII? Consistent with previous studies we have shown that SodCII is proteolytically degraded in the phagosome and/or phagolysosome in Perforin-2 proficient cells (Krishnakumar et al., 2004; Krishnakumar et al., 2007). However, the degradation of SodCII is significantly attenuated in phagocytes lacking Perforin-2. Similar results were observed with cytoplasmic DnaK in BMDM. This was not due to a general defect in protease activity in the phagolysosome because the surface antigen flagellin was degraded whether or not Perforin-2 was present. Thus, Perforin-2 facilitates the degradation of antigens contained within the envelope of phagocytosed bacteria. However, it is unlikely that Perforin-2 is itself a protease because it lacks significant homology to a known protease or protease motif. What Perforin-2 does have is a MACPF domain (Figure 7) (McCormack and Podack, 2015; Ni and Gilbert, 2017; Podack and Munson, 2016). This suggest that Perforin-2 is a pore forming protein and putative Perforin-2 pores have been imaged by transmission electron microscropy (McCormack et al., 2015a). However to date it has not been possible to confirm that the imaged structures contain Perforin-2 due to the absence of suitable antibodies. Nor has it been confirmed that the structures are in fact pores. However, other MACPF containing proteins such as complement protein C9 and Perforin have been shown to polymerize and form pores in lipid membranes (Dudkina et al., 2016; Law et al., 2010; Podack et al., 1989; Podack et al., 1982; Podack and Tschopp, 1982). For example, 22 monomers of C9 polymerize to form a pore in the outer membrane of gram-negative bacteria with an inner diameter of 120 Å (Dudkina et al., 2016; Podack et al., 1982; Tschopp and Podack, 1981). Because Perforin-2 is a type I transmembrane protein with its membrane spanning alpha helix near its carboxy-terminus, the MACPF domain of Perforin-2 would reside in the lumen of endosomes and phagosomes. In this orientation its MACPF domain would reside in the same compartment as phagocytosed bacteria. Thus, we propose that Perforin-2 polymerizes – perhaps after cleavage from its transmembrane domain by a lysosomal protease– and forms pores in the envelope of phagocytosed bacteria (Figure 7). This model is consistent with our experimental observations with *S. typhimurium* since the protease that degrades SodCII would enter the periplasmic space through poly-Perforin-2 pores. Because SodCII is not anchored in the periplasm, it may also diffuse through the pore and be degraded in the lumen of the phagosome (Kim et al., 2010; Krishnakumar et al., 2004; Krishnakumar et al., 2007). In either case SodCII is able to persist in the periplasm and protect the bacterium from the bactericidal effects of ROS when Perforin-2 is absent.

**Figure 7.**
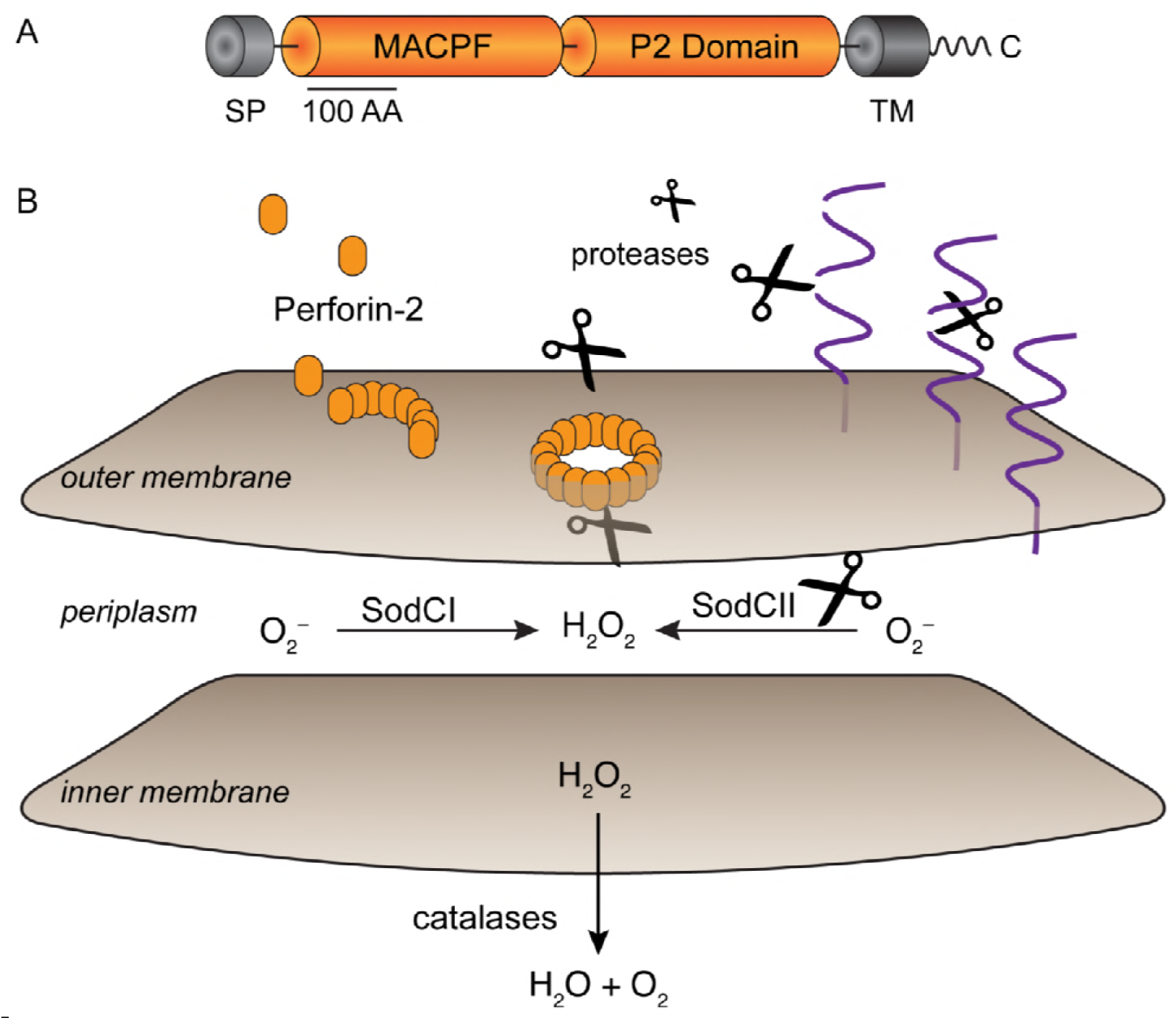
Model for Perforin-2 dependent degradation of SodCII. (A)Domain organization of Perforin-2. MACPF domains are present in other proteins of the innate immune system and in the case of complement protein C9 and Perforin-1 have been shown to polymerize and form pores in lipid membranes. The P2 domain is a domain of unknown function that is conserved in orthologs of Perforin-2. As a type I transmembrane protein the MACPF domain of Perforin-2 would reside within the lumen of endosomes. (B)Upon activation and likely cleavage from its transmembrane domain Perforin-2 polymerizes on the outer membrane of *S. typhimurium*. Subsequent pore formation allows proteases –and other antimicrobial effectors– to enter the periplasmic space and degrade SodCII. Alternatively, SodCII may diffuse through the poly-Perforin-2 pore and be destroyed in the lumen of the phagolysosome. Pores do not lead to the inactivation of SodCI because it is protease resistant and tethered within the periplasm. Abbreviations; SP, signal peptide; TM, transmembrane domain

In addition to Perforin-2 there is evidence that the cathelicidins CRAMP and LL-37 also play a role in disrupting the envelope of phagocytosed gram-negative bacteria in murine and human macrophages respectively (Kim et al., 2010; Rosenberger et al., 2004; Sonawane et al., 2011; Stephan et al., 2016). Of particular relevance to this study are previous studies that have shown that CRAMP is active against phagocytosed *S. typhimurium* (Kim et al., 2010; Rosenberger et al., 2004). In one it was shown that CRAMP inhibits the division of *S. typhimurium.* The result was filamentous bacteria and it was further shown that filamentation was protease dependent (Rosenberger et al., 2004). In another study it was shown that CRAMP is associated with the loss of SodCII from the periplasm of *S. typhimurium* in vitro (Kim et al., 2010). Moreover SodCII promoted the survival of *S. typhimurium* in CRAMP deficient but not proficient mice. However in the latter study the authors also noted that the loss of CRAMP failed to fully abolish the degradation of SodCII. Our study suggest that this is most likely due to the activity of Perforin-2. Although it is clear that Perforin-2 and cathelicidins can act independently of one another, it remains to be determined if they also act synergistically. In the case of gram-negative bacteria a particularly intriguing model is the possibility that Perforin-2 forms a conduit in the outer membrane through which cathelicidin or other antimicrobial peptides transit to reach the bacterial inner membrane. Alternatively the deployment of independent mechanisms to disrupt the envelope of phagocytosed bacteria may be an insurance strategy against pathogen resistance to any particular mechanism.

## Material And Methods

### Mice

Perforin-2 -/- 129X1/SvJ mice were produced at the University of Miami Miller School of Medicine Transgenic Core Facility as previously described (McCormack et al., 2015b). Wild-type 129X1/SvJ and Perforin-2 -/- mice of either sex were used at 2 to 6 months of age. Mice received food and water *ad libitum* and were housed at an ambient temperature of 23°C on a 12 h light/dark cycle under specific pathogen-free conditions. Mice were euthanized by CO_2_ inhalation followed by cervical dislocation. All procedures with animals were reviewed and approved by the University of Miami’s Institutional Animal Care and Use Committee.

### Strains and Plasmids

Bacterial strains are listed in Table 1. Primer sequences are listed in Table 2. Strains constructed for this study are isogenic derivatives of *S*. *typhimurium* strain LT2 (Nikaido et al., 1967). Plasmid pKD4 (Datsenko and Wanner, 2000) was used in PCR with primer pairs sodCI-P1/sodC1-P2 or sodCII-P1/sodCII-P2 to generate kanamycin resistance cassettes bracketed by recognition sites for FLP recombinase and flanking sequences targeting *sodCI* or *sodCII*. Deletions of *sodCI* or *sodCII* were generated by λRed-mediated recombination of the cassettes into LT2 as described (Datsenko and Wanner, 2000). Recombinants were selected on LB agar plates with kanamycin. Recombination sites were verified by PCR with flanking primer pairs sodCI-MfeI/sodCI-HindIII for *sodCI::kan* and sodCII-EcoRI/sodCII-HindIII for *sodCII::kan*. For the construction of double deletions FLP recombinase was used to excise the first cassette prior to insertion of the second.

**Table 1.**
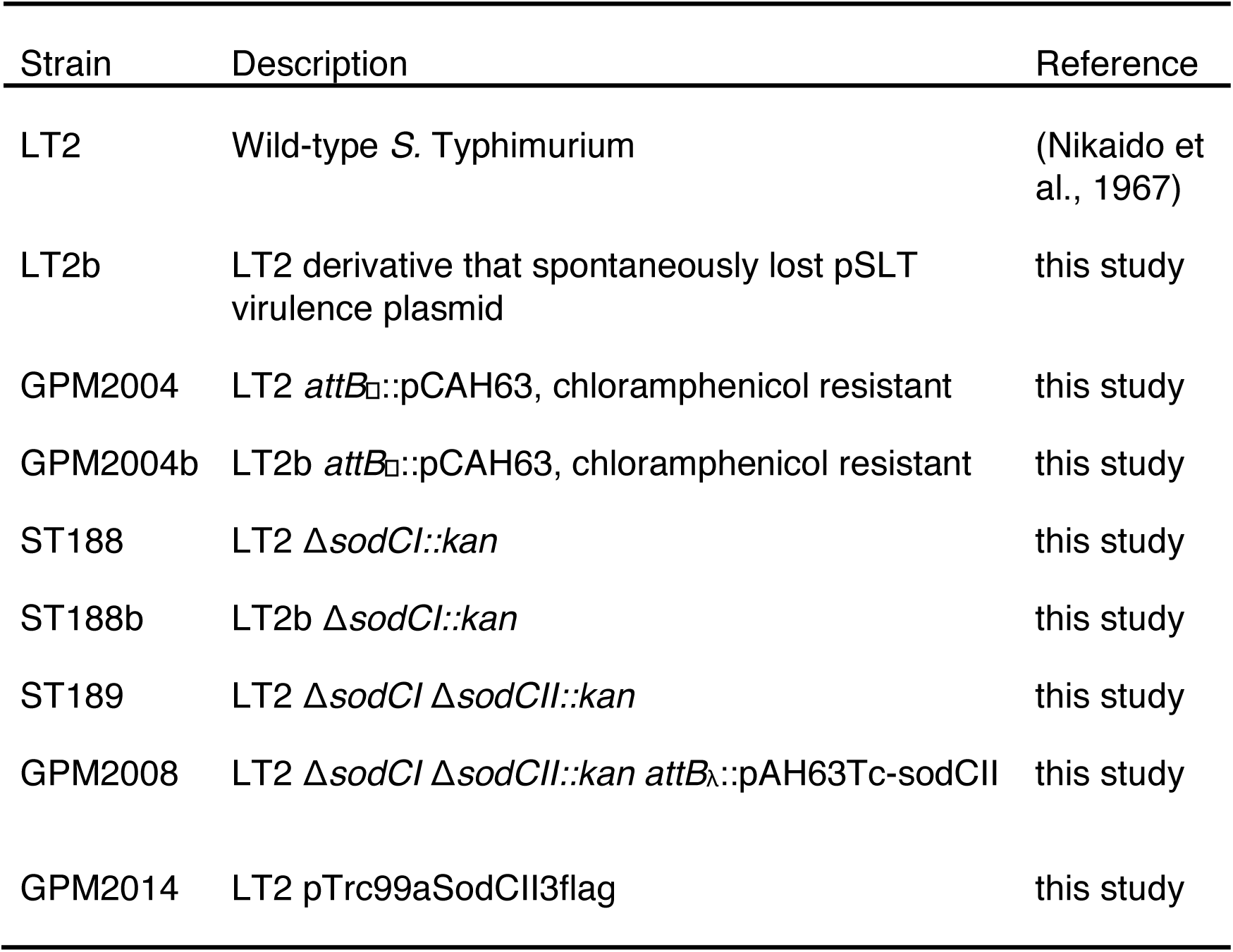
Bacteria strains and plasmids

| Strain | Description | Reference |
| --- | --- | --- |
| LT2 | Wild-type *S. Typhimurium* | (Nikaido et al., 1967) |
| LT2b | LT2 derivative that spontaneously lost pSLT virulence plasmid | this study |
| GPM2004 | LT2 *attB*::pCAH63, chloramphenicol resistant | this study |
| GPM2004b | LT2b *attB*::pCAH63, chloramphenicol resistant | this study |
| ST188 | LT2 Δ*sodCI*::*kan* | this study |
| ST188b | LT2b Δ*sodCI*::*kan* | this study |
| ST189 | LT2 Δ*sodCI* Δ*sodCII*::*kan* | this study |
| GPM2008 | LT2 Δ*sodCI* Δ*sodCII*::*kan attB*::pAH63Tc-sodCII | this study |
| GPM2014 | LT2 pTrc99aSodCII3flag | this study |

**Table 2.**
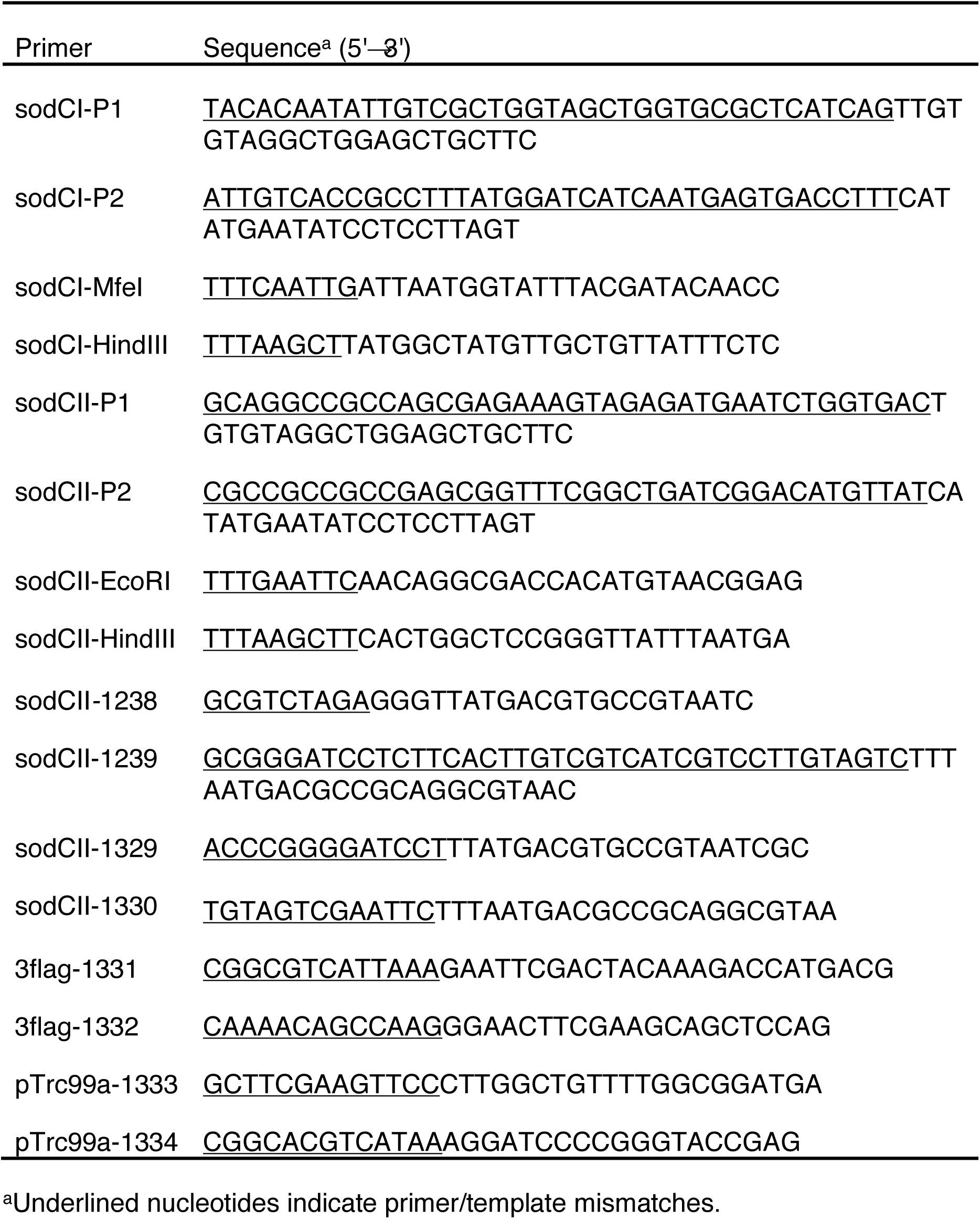
Oligonucleotide Primers

The *sodCII* gene was cloned from LT2 by PCR with primers sodCII-1238/sodCII-1239. The PCR product was digested with XbaI and BamHI then ligated into the same sites of pAH63Tc, a derivative of pAH63 (Haldimann and Wanner, 2001) in which the kanamycin resistance gene was replaced with one conferring resistance to tetracycline. To complement *sodCII* mutants the resulting plasmid, pAH63Tc-sodCII, was integrated into *attB*_λ_ of relevant strains by a site specific recombinase as previously described (Haldimann and Wanner, 2001). The chloramphenicol resistant strain GPM2004 was constructed by integration of pCAH63 (Haldimann and Wanner, 2001) into *attB*_λ_ of LT2. Plasmid pTrc99aSodCII3flag was constructed by HiFi DNA assembly (New England Biolabs) of three PCR products: one carrying *sodCII* amplified from pAH63Tc- sodCII with primers sodCII-1329/sodCII-1330, another carrying a triple FLAG epitope amplified from pSUB11 (Uzzau et al., 2001) with primers 3flag-1331/3flag-1332, and the vector backbone amplfied from pTrc99a (Amann et al., 1988) with primers pTrc99a-1333/pTrc99a-1334. Bacteria were cultured in Luria-Bertani (LB) broth or plates. As appropriate antibiotics were used at the following concentrations: chloramphenicol, 10 µg/ml; ampicillin, 100 µg/ml; kanamycin, 50 µg/ml; tetracycline 7.5 µg/ml.

### Cell preparation

Peritoneal exudate macrophages (PEMs) were isolated as previously described (Zhang et al., 2008). Briefly, mice were injected intraperitoneally with 1 ml of 4% Brewer thioglycollate medium and peritoneal exudates were recovered after 4 days. A modified schedule was used to collect PEM for ROS assays (Nathan and Root, 1977). In brief, mice were inoculated on days 1 and 7, and exudates were collected on day 14. Resting macrophages were collected from the peritoneal cavities of untreated mice. Peritoneal neutrophils were elicited and recovered as previously described (Luo and Dorf, 2001). Briefly mice were injected intraperitoneally with 1 ml 9 % casein in PBS (2.7 mM KCl, 4.3 mM Na_2_HPO_4_, 1.47 mM KH_2_PO_4_, 137 mM NaCl, pH 7.4) containing 0.9 mM CaCl_2_ and 0.5 mM MgCl_2_. A second injection was administered 16 h after the first and exudates collected 3 h later. Cells were maintained in IMDM (Gibco) supplemented with 10 % fetal bovine serum (FBS) (Gibco) at 37°C in 5% CO2. Bone marrow derived macrophage (BMDM) were isolated from the femurs from the Perforin-2 WT and KO mice. The cells were cultured in IMDM medium supplemented with 10% FBS, 20% L929 conditioned medium, and 10 mM L-glutamine. BMDM were cultured for 6 days before use, and the culture medium was changed every 2 days.

### Intracellular killing assays

Intracellular gentamicin protection assays were performed as previously described (Laroux et al., 2005; Lutwyche et al., 1998; McCormack et al., 2013). Briefly, 3 × 10^5^ PEM were seeded in 24 well plates in IMDM, 10 % FBS and incubated at 37°C in 5% CO_2_. Murine IFN-γ was added 14 hours before infection at a final concentration of 50 ng/ml. Overnight cultures of bacteria were diluted 33 fold in LB and cultured aerobically at 37°C for 3 hpurs to mid-log at which point the optical absorbance of the culture at 600nm was ca. 0.6. Bacteria were added at a multiplicity of infection (MOI) between 20 to 50 and plates incubated for 45 to 60 min to allow for uptake of bacteria. Cells were then washed three times with PBS and fresh culture medium containing 50 µg/ml gentamicin was added to each well to kill extracellular bacteria. After 2 hours the concentration of gentamicin was reduced to 5 µg/ml. At selected time points gentamicin was removed by PBS washes and bacteria were recovered by lysis of mammalian cells in strerile water with 0.1% Triton X-100. Lysates were serially diluted in PBS and the bacteria were enumerated on LB agar plates.

### ROS detection

3 × 10^5^ PEM were seeded in 96-well opaque white plates in 200 µl of phenol red– free IMDM, 10% FBS and primed for 14 hours with murine IFN-γ (Biolegend) at a final concentration of 50 ng/ml. Alternatively, 3 × 10^5^ neutrophils were seeded in KRPG buffer (145 mM NaCl, 4.86 mM KCl, 5.7 mM sodium phosphate, 0.54 mM CaCl_2_, 1.22 mM MgSO_4_, 5.5 mM Glucose, pH 7.35). For ROS production in neutrophils, cells were incubated with 1 mM luminol (Sigma) for 3 min. before adding phorbol myristate acetate (PMA)(InvivoGen) or lipopolysaccharides (LPS) (InvivoGen) to a final concentration of 100 ng/ml. Luminescence was read in an EnVision (PerkinElmer) plate reader. Some wells were treated with 10 nM diphenyleneiodonium chloride (DPI) (Sigma) –a NAPDH oxidase inhibitor– for 30 min prior to addition of luminol. The same procedures were used with macrophages except the luminol enhancer Diogenes (National Diagnostics) was also used (Yamazaki et al., 2011).

### Recovery of phagocytosed bacteria and immunodetection

*S. typhimurium* strain GPM2014 */* pTrc99aSodCII3flag was cultured aerobically overnight in LB with ampicillin at 37°C. The bacteria were pelleted, washed with sterile PBS and aliqouts of the inoculum were frozen at -80 °C for later analysis. Aliquots of the inoculum were also used to infect wild-type and Perforin-2 -/- PEMs at a multiplicity of infection of about 20-50. After 1 hour incubation, the cells were washed three times with PBS then IMDM with 10% FBS and 50 µg/ml gentamicin was added to each well. This initial addition of gentamicin marked the 0 hour time point. After 2 hours the concentratio of gentamicin was decreased to 5 µg/ml. After 16 hours the cells were washed with PBS three times, then PBS, 0.1% Triton-X 100 containing a proteinase inhibitor cocktail (Roche, Basel, Switzerland) was added to each well. After a 5 min. incubation at 37°C the cells were manually detached with a cell scraper, the bacterial cells were harvested by centrifuged at 4°C at 10, 000 *g* × 10 min, and the supernatant was removed. Recovered phagocytosed bacteria (approximate 10^6^CFU) and bacteria from the original LB culture were boiled in Laemmli loading buffer for 7 min. The same infection procedure was conducted with wild-type and Perforin-2 -/- BMDMs except the phagocytosed bacteria were collected at 1, 3 and 6 hours. The protein samples were separated on 4-20% gradient SDS-PAGE gels and transferred to nitrocellulose membranes. The membranes were blocked with 5% non-fat milk in Tris buffered saline containing 0.1% Tween-20 (TBST) for 2 hours and then incubated at 4°C for 16 hours with primary antibodies anti-FliC (1:1000, Invivogen), anti-Flag (1:5000, Sigma) or anti-DnaK (1:5000, Abcam) diluted in TBST with 5% non-fat milk. After three washes with TBST the membranes were incubated at 37°C for 1 hour with anti-mouse horseradish peroxidase-labeled secondary antibody (Jackson ImmunoResearch Laboratories, West Grove, PA, USA) diluted 1:5000 in TBST with 5% non-fat milk. An Odyssey FC Imaging System (LI-COR, Lincoln, NE, United States) was used to detect and quantify chemilunescence after addition of SuperSignal West Pico Chemiluminescent Substrate (Thermo Fisher Scientific).

### Murine infections

Bacteria were cultured overnight in LB medium at 37°C and diluted in sterile PBS. For competition assays selected strains of *S. typhimurium* were mixed at a 1:1 ratio. Perforin-2 +/+ and -/- 129X1/SvJ mice were inoculated by intraperitoneal (i.p.) injection. The CFU of each inoculum was quantified by plating and ranged from 500 to 1,000 total CFUs. Spleens and livers were collected four days after inoculation, and then homogenized (Omni International) in 500 µl PBST (PBS, 0.1% Tween-20). Homogenates were diluted in sterile PBS and plated in triplicate on LB agar with antibiotic selection as appropriate for each strain in the initial inoculum. For each spleen and liver, mean CFUs were used to calculate competitive indices (CI) according to the following formula: CI = (strain A recovered / strain B recovered) / (strain A inoculum / strain B inoculum).

### Statistical analysis

Statistical analysis was performed with GraphPad Prism 7 software. Data represent the mean ± standard deviation (SD). Statistical difference was determined by the Student’s *t*-test. *P* values ≤ 0.05 were considered statistically significant. The number of independent experimental replicates is indicated by *n*.

## Acknowledgments

Research reported in this publication was supported by Grant Number AI110810 from the National Institute of Allergy and Infectious Diseases of the National Institutes of Health (NIAID NIH). Its contents are solely the responsibility of the authors and do not necessarily represent the official views of the NIH.

## Author Contributions

Conceptualization, F.B., R.M.M., M.G.L., and G.P.M.; Methodology, F.B., R.M.M., and G.P.M.; Investigation, F.B.; Validation, F.B and G.P.M.; Formal Analysis, F.B.; Resources, S.H., G.V.P., and R.M.M.; Writing – Original Draft, F.B. and G.P.M.; Writing – Review & Editing, F.B., R.M.M., S.H., G.V.P., M.G.L., and G.P.M.; Visualization, F.B. and G.P.M.; Supervision, M.G.L. and G.P.M.; Funding Acquistion, G.P.M.

## Declaration of Interests

The authors declare no competing interests.

